# High Resolution Single Particle Cryo-Electron Microscopy using Beam-Image Shift

**DOI:** 10.1101/306241

**Authors:** Anchi Cheng, Edward T. Eng, Lambertus Alink, William J. Rice, Kelsey D. Jordan, Laura Y. Kim, Clinton S. Potter, Bridget Carragher

## Abstract

Automated data acquisition is now used widely for the single-particle averaging approach to reconstruction of three-dimensional (3D) volumes of biological complexes preserved in vitreous ice and imaged in a transmission electron microscope (cryo-EM). Automation has become integral to this method because of the very large number of particle images required to be averaged in order to overcome the typically low signal-to-noise ratio of these images.

For optimal efficiency, all automated data acquisition software packages employ some degree of beam-image shift because this method is fast and accurate (+/− 0.1 μm). Relocation to a targeted area under low-dose conditions can only be achieved using stage movements in combination with multiple iterations or long relaxation times, both reducing efficiency. It is, however, well known that applying beam-image shift induces beam-tilt and hence structure phase error. A π/4 phase error is considered as the worst that can be accepted, and is used as an argument against the use of any beam-image shift for high resolution data collection.

In this study, we performed cryo-EM single-particle reconstructions on a T20S proteasome sample using applied beam-image shifts corresponding to beam tilts from 0 to 10 mrad. To evaluate the results we compared the FSC values, and examined the water density peaks in the 3D map. We conclude that the π/4 phase error does not limit the validity of the 3D reconstruction from single-particle averaging beyond the π/4 resolution limit.

## Introduction

The single particle averaging approach to reconstruct three-dimensional (3D) volumes of biological complexes preserved in vitreous ice and imaged by cryo electron microscopy (cryo-EM) is a powerful technique with increasing numbers of structures reconstructed to a resolution range of 3 Å or better (Crowther, 2016). There were 25 sub-3 Å maps (arising from 10 published papers) deposited in the EMDB in the last six months. Due to the radiation sensitivity of biological samples preserved in vitreous ice, to achieve high resolutions, very large numbers of particle images have to be acquired, each at a previously unexposed area. This makes the acquisition process slow, and, before automated data acquisition was developed (Tan et al., 2016), it was also very labor intensive.

Automated data acquisition is typically performed with a series of targeting and imaging steps at progressively higher magnifications. This workflow improves targeting accuracy while practicing low-dose methods, which is critical for high resolution cryo-EM. Automated eucentric height adjustment and object lens focusing are inserted between some of the steps. To image at the targets selected at lower magnifications, either a stage movement or image deflector with a compensated beam shift is used to center the targeted image on the detector.

All popular automated data acquisition software packages used today (Leginon (Suloway et al., 2005), SerialEM (Mastronarde, 2005) and EPU) have this capability and often employ some degree of beam-image shift (as opposed to stage shift) to optimize the efficiency of the workflow for high magnification targeting. This is because even the best mechanical stages on the most commonly used electron microscopes can only achieve a reproducibility of +/− 1 μm when requested to move to a target some distance away (e.g. 20 μm) (Cheng et al., 2016). Alternatively, the high precision obtained by some mechanical or piezo-driven stages requires long relaxation times to reduce residual stage drift. We have observed, over numerous projects, that for final high magnification targeting, a moderate beam-image shift (~2 μm) can be used and will produce a map of very high quality (even as high at 2.5 Å) in half the time required when using stage shift.

Despite this success, using beam-image shift is against the conventional wisdom that this approach will induce beam tilt, and produce distortion in the image by changing the phase of the structure factor. Here we present a series of experiments exploring the limits of the resolution that can be achieved for SP reconstruction when beam-image shifts corresponding to beam tilts from 0-10mrad are used for targeting. The results are much better than might be expected and we provide a possible explanation for why this may be so.

## Theory

Lens aberrations distort the electron wave. As we seek higher resolution, more of these aberrations have to be considered. Defocus, and 2-fold astigmatism are commonly corrected for cryo-EM images of biological molecules. A further important aberration that needs to be considered, at least when the spherical aberration constant, *Cs*, is non-zero, is the coma effect produced from beam tilt, *θ* (Glaeser et al., 2011; Uhlemann and Haider, 1998).

The coma effect causes both a defocus change in the direction of the beam tilt as well as an azimuthally varying phase error of the structure factor, *Δφ*:

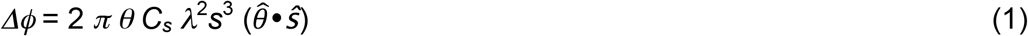

where *λ* is the electron wavelength, *s* the spatial frequency, 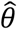 is a unit vector in the direction of the beam tilt and *ŝ* is a unit vector in the direction of the spatial frequency vector. This structure factor phase error creates a distortion of the image that is not correctable without knowing the magnitude and direction of the beam tilt. As discussed in Glaeser et al. (2011), the “worst that one could accept” phase error is defined as π/4. Applying that value in Equation 1, for a known beam tilt, the π/4 resolution limit is:

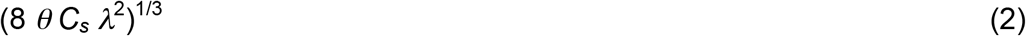

Table 1 lists a few beam tilts relevant to this manuscript and their π/4 and π/2 resolution limits.

**Table 1:** Phase error resolution limit at 300kV and microscope with 2.7 mm Cs in comparison to the experimental (Expt) FSC_0.143_

| Beam Tilt (mrad) | 0.25 | 0.33 | 0.5 | 1.00 | 1.30 | 10.0 |
| --- | --- | --- | --- | --- | --- | --- |
| $\pi/4$ resolution limit (Å) | 2.75 | 3.0 | 3.47 | 4.36 | 4.77 | 9.41 |
| $\pi/2$ resolution limit (Å) | 2.19 | 2.40 | 2.75 | 3.47 | 3.79 | 7.48 |
| +tilt expt $FSC_{0.143}$ resolution (Å) | | | 3.05 | | 3.39 | 5.37 |
| +/-tilt expt $FSC_{0.143}$ resolution (Å) | | 2.89 | 2.92 | | 3.16 | |

According to this calculation, even though a single image taken at a given beam tilt may appear to have features resolved beyond the π/4 resolution limit, the distortion of the feature will be considered as not representing the structure.

Figure 1 illustrates this distortion on a phantom images consisting of two white squares separated by their length (Figure 1a). If the center of the object is determined by an algorithm, the positions of these centers will be found half way between the squares, as indicated by the black line throughout Fig 1. If a severe simulated coma effect is applied according to Equation 1, the objects are no longer symmetrical around their original centers (Figure 1b, 1c).

**Figure 1:**
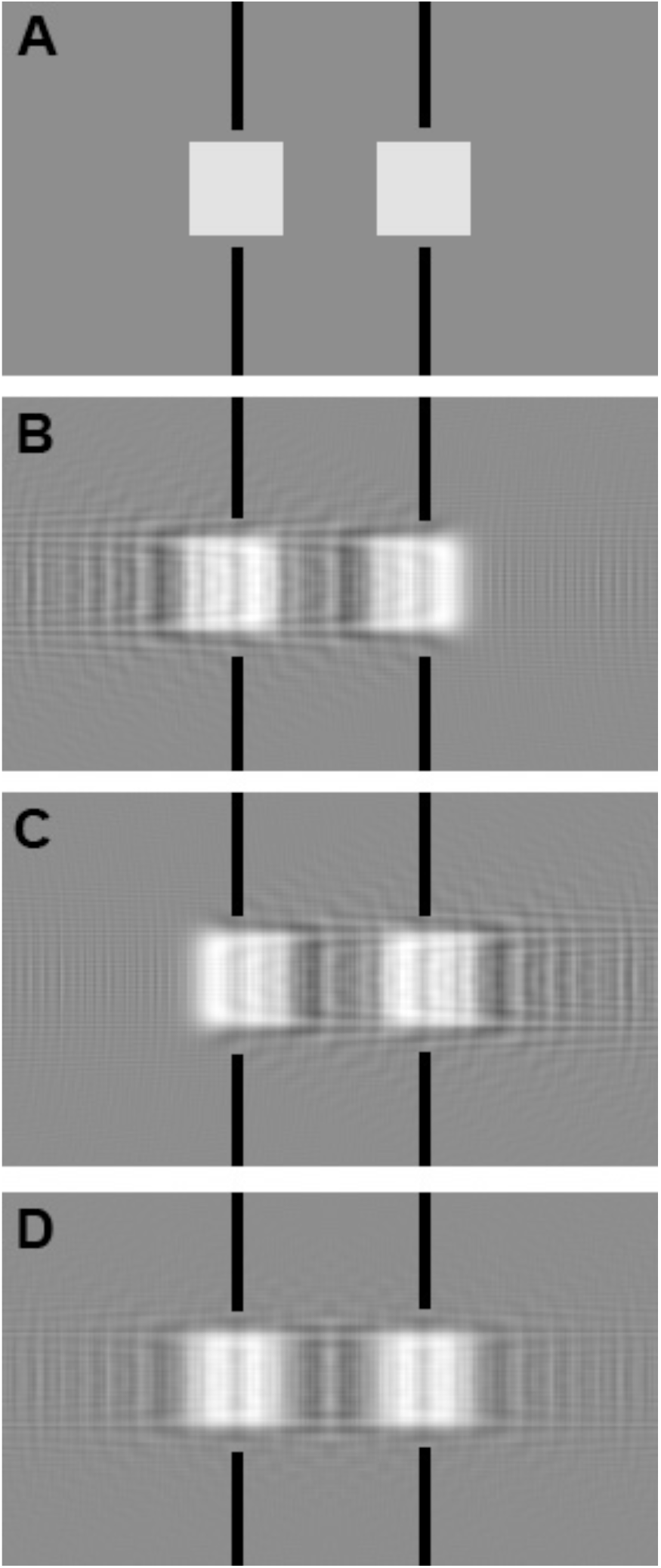
Simulation of coma effect on image structure. (A) phantom image; (B) and (C) simulated beam tilt in the horizontal plane toward right and left, respectively; (D) averaging of (B) and (C) produces an image centered at the same position as the input.

We observe that *Δφ* becomes −*Δφ* if 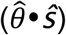 changes sign. This situation may happen either because the beam is tilted to −***θ***, or the object is rotated by 180 ° in-plane. While the distortion of the features are not the same, the vector addition of the structure factor imposed during the averaging process cancels the phase errors but also reduces the amplitudes. This effect means that the accuracy of the feature location is not affected by the coma at any resolution when we combine data from two opposite tilts, represented as +/−*θ* throughout this manuscript. This canceling effect on coma is illustrated in Figure 1. Figure 1B and 1C show the effects of coma in opposite directions, leading to different distortions. When the two images are averaged, the objects are again centered as shown in Figure 1D. The averaged image has blurred edges since the higher resolution structure factors are weighted down by a factor of cos *Δφ*.

Single-particle averaging collects large numbers of projection images of the same structure in random orientations and then computationally aligns and averages them to reconstruct the 3D map. Considering these large numbers of input images, it is reasonable to assume that there are always pairs of particle images that are at the same 3D orientation with one experiencing beam tilt at +*θ* while the other beam tilt is at −*θ*. Therefore, until the single-sideband phase error is π/2, the combined coma-effect is an *s* dependent amplitude reduction of the structure factor without phase errors. Table 1 also lists the π/2 resolution limit for various beam tilts.

## Material and Methods

Four experiments using different estimated beam tilts were analyzed in this study: (1) Typical +/−0.33 mrad; (2) 0, +/−0.5, +/−1.33 mrad; (3) 5 mrad; (4) 10 mrad. Data for each of the four experiments was acquired on different grids. Supplemental Table 1S summaries the experiment design.

### Sample

*Thermoplasma acidophilum* 20S proteasome was a gift from Yifan Cheng, Znlin Yu, and Kiyoshi Egami. The received stock was separated into small aliquots and stored at −80 °C.

### Grid Preparation

3μl of freshly thawed protein was applied to plasma-cleaned C-flat 1.2/1/3 400 mesh Cu holey carbon grids (Protochips, Raleigh, North Carolina), blotted for 2.5 s after a 30 s wait time, and then plunge frozen in liquid ethane, cooled by liquid nitrogen, using the Cryoplunge 3 (Gatan) at 75% relative humidity.

### Microcopy

Thermo Fisher Scientific Titan Krios operated at 300 kV and Gatan K2 Summit camera in counting mode were used in all experiments described. The Cs value is 2.7 mm. The experiments shown are from two different microscopes differing only in that that one has a Gatan BioQuantum energy filter; we did not observe any trends associated with energy filtering. Pixel size was either 1.06 or 1.10 Å. A 100 μm objective aperture was used except in the 5 and 10 mrad beam tilt experiment (Experiment 3 and 4).

### Imaging

Movies in the four experiments were collected in counting mode using Leginon (Suloway et al., 2005) at a dose rate of 8.0 e-/Å^2^/s with a total exposure time of 5-7 seconds, for an accumulated dose of 35 – 42 e^−^/Å^2^. Intermediate frames were recorded every 0.1 or 0.2 seconds for a total of 25 – 60 frames per micrograph. Defocus values range from approximately 1.0 – 3.0 μm.

### Beam tilt induced by beam-image shift

For a well-aligned microscope with minimal induced beam tilt, the measurement was performed according to the procedure previously described (Cheng et al., 2016). For experiments using an intentionally misaligned beam (Experiment 2, 3, and 4), beam tilt was estimated from observing the movement of the diffraction ring from a grating replica gold-palladium cluster relative to the 70 μm objective aperture. The (111) and (200) diffraction ring scatters at 8.52 and 9.84 mrad at 300 kV high-tension. The radius difference between these two rings in the diffraction image therefore provides the scale calibration.

### Experiment 1; Typical +/−0.33 mrad beam tilt imaging

The typical imaging condition we use with a well-aligned scope introduces up to 0.33 mrad when 4 targets are selected in the lower-magnification overview image requiring an image shift distance of ~1.7 μm to center the target at the high magnification setting (Figure 2A). Axial-coma was minimized with the help of Zemlin tableau.

**Figure 2.**
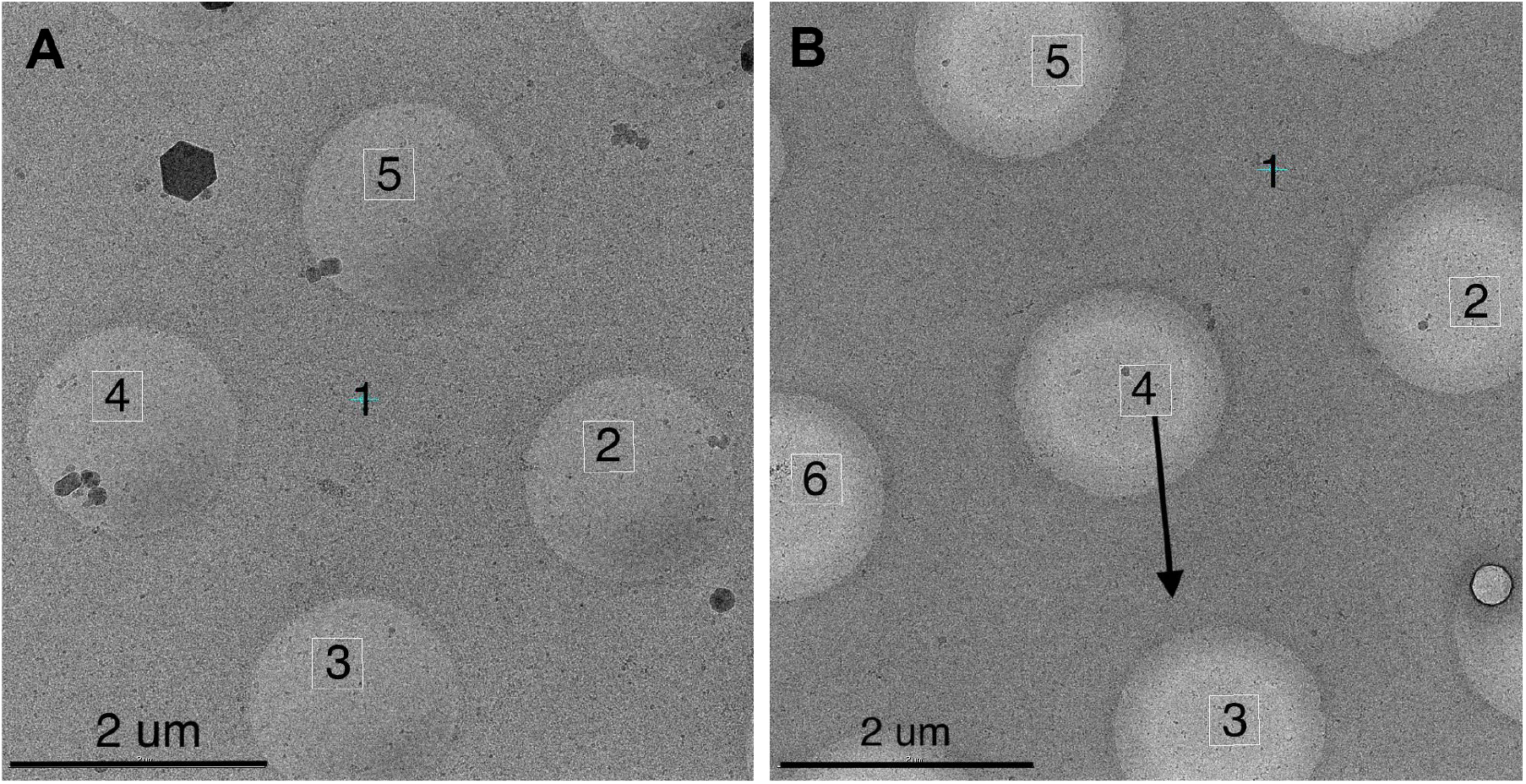
Targeting scheme. The numbers represent the order of target processing, Target 1 is where autofocusing is performed. (A) Standard 1.7 μm beam-image shift targeting used in all Experiments except 2. Targets are determined automatically to adjust with the hole location variation from stage movement inaccuracy. (B) 2.6 μm beam-image shift targeting used in Experiment 2 for up to 1.3 mrad beam tilt imaging. A fixed pattern of targets was used for all imaging. The black arrow is roughly aligned to the image shift x direction and was the direction along which the beam-shift and beam-tilt pivot points were misaligned. Target 4 was chosen to require no beam-image shift. Target 3 and 5 have the maximal beam tilt at this orientation while target 2 and 6 have minimal beam tilt.

### Experiment 2; 0, +/−0.5 and +/−1.3 mrad beam tilt imaging

To induce larger than normal beam tilts at given beam-image shifts, we modified the beam shift and beam tilt pivot point x in the registry as well as beam-image calibrations. Beam shift and beam tilt pivot points are generally aligned around zero beam-tilt so that applied beam shift receives balanced deflector currents to minimize the change of beam tilt from zero. We changed the scale of the correction the deflectors make so that it loses the balance in the x direction. Beam-image calibration calibrates the scale and direction of beam shift required to keep the beam stationary on the detector at a given change of image shift deflector current. The modification of the pivot point makes the original calibration invalid. Therefore a round of beam-image calibration was performed. The amount of beam tilt induced in beam-image shift of 2.6 μm, the distance of our imaging targets from the center, was then estimated as described above. Leginon was set up with 5 targets in the lower-magnification overview image as shown in Figure 2B. The center target acquires a high-magnification image with no beam image shift applied. Beam-image shifts were required to image the surrounding more distant targets. Two of these targets have maximal beam tilt and the other two have minimal beam tilt as indicated in Figure 2B.

### Experiment 3 and 4; 5 and 10 mrad beam tilt imaging, respectively

It was not possible to induce beam tilts larger than 1.3 mrad without significant targeting error in the beam-image shift. The 5 mrad beam tilt was induced by applying a constant beam tilt value in a custom Leginon node only during final high-magnification image acquisition. The four target beam-imaging shift protocol described in Figure 2A was used. The objective aperture was removed to minimize side-band effects at these large beam tilts. The 10 mrad beam tilt imaging was performed with the same protocol.

### Image Processing

Movies recorded on the K2 were aligned using MotionCor2 with dose weighting (Zheng et al., 2017) and CTF estimation was performed with CTFFIND4 (Rohou and Grigorieff, 2015). For the first 50 images particles were picked automatically using the Appion DoG Picker (Voss et al., 2009), extracted, and subjected to 2D classification in RELION (Scheres, 2012) to create templates for another round of particle picking using FindEM (Roseman, 2004). The picked particles were extracted and subjected to 2D classification and the best classes was subjected to ab initio reconstruction to create an initial model in cryoSPARC (Punjani et al., 2017). This model was used for 3D classification in RELION or heterogeneous refinement in cryoSPARC. For the final reconstruction, particles corresponding to the best two 3D classes were selected and subjected to 3D refinement in cryoSPARC using a box size of 256×256 pixels. An earlier experiment (Experiment 1) with its micrographs aligned using MotionCorr (Li et al., 2013) also underwent particle polishing in RELION.

3D reconstructions were soft masked and sharpened with auto-B-factor determination using cryoSPARC or RELION. The program applied (2*FSC/(FSC+1))^1/2^ weighting as in (Rosenthal and Henderson, 2003) and fitting of a straight line through the Guinier plot to find the B-factor.

## Results

We collected and analyzed particle images taken at various beam tilts. Figure 3 summarizes the FSC_0.143_ resolution of each of the resulting 3D maps defined in RELION (Scheres, 2012) and cryoSPARC (Punjani et al., 2017). All beam-tilted data sets achieved resolutions higher than the π/4 limit and those tilted more than 1.0 mrad even exceeded the π/2 limit. It is thus clear that the coma effect phase error does not limit the FSC_0.143_ resolution.

**Figure 3:**
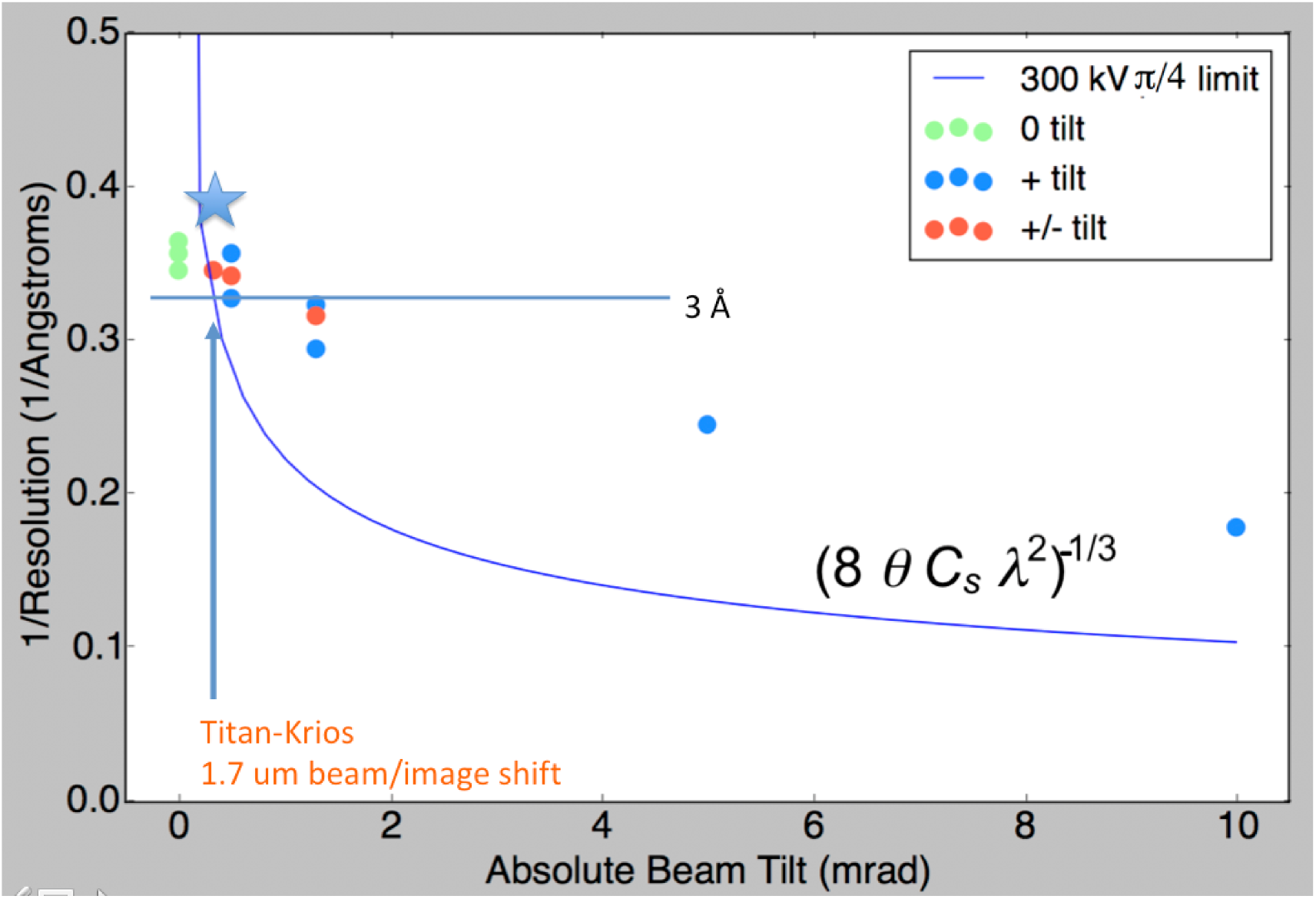
The FSC_0.143_ resolution achieved from maps reconstructed using images with defined beam tilt applied. The phase error resolution limits at π/4 is shown as theoretical references. The star marks the highest resolution proteasome structure obtained with 1.7 μm beam/image shift protocol, although at a smaller pixel size.

We also examined water density peaks identified in the 2.8 Å structure (EMD-6287 and deposited in PDB as 6BDF) to validate both the existence and the position of these features since they are only separable at better than 3 Å resolution and are more sensitive to phase error of the structure factor at higher resolution. Figure 4 shows one such region. The two water peaks we tracked in various maps weaken as the reported FSC_0.143_ resolution decreased while the center of the peak is well-preserved. More water density tracking is presented in Supplementary Material.

**Figure 4:**
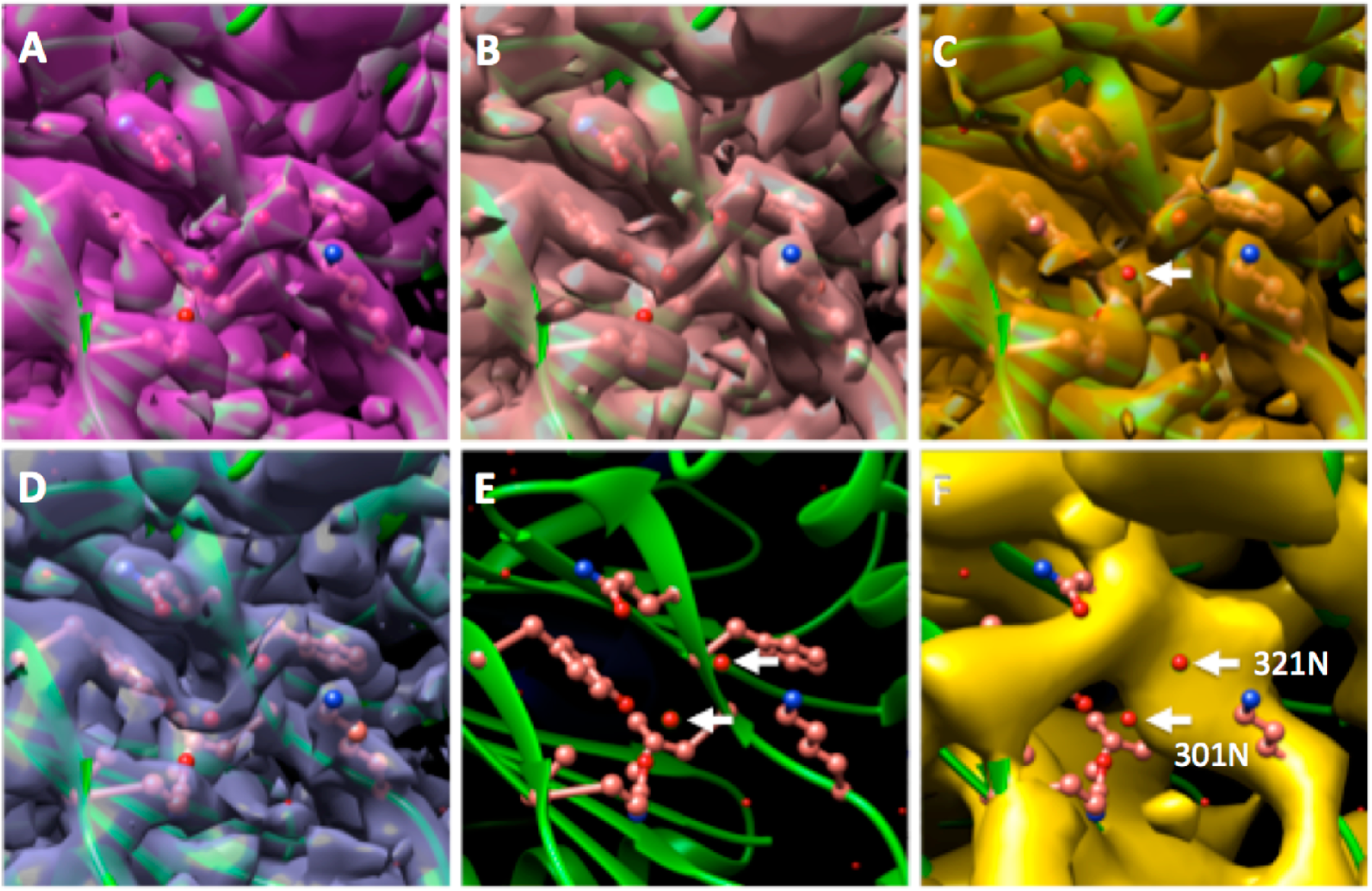
Water density as validation of the correct phase of the density maps. Panels A,B,C are from the same EM grid and same experimental session with different beam tilts (Experiment 2). (D) is from EMD-6287, (E) shows the PDB modeled from (D) that is placed in all maps. The estimated beam tilts are 0, +/− 0.5, +/−1.33, 0, +10 mrad for panel A, B, C, D, F, respectively. The white arrows in (E) and (F) shows the water molecule we track in the various maps. The arrow in (C) points to the 301N water that was not represented by density at the contour level. Maps were aligned against EMD-6287 with fit-in-map function in UCSF Chimera (Pettersen et al., 2004). Surface contours were thresholded such that 1% of voxels lie above the threshold level.

## Discussion

The capital equipment and operational cost of high-end cryo-EM microscopes and the competitiveness of the structural biology field demands that these instruments are used as efficiently as possible. Given that mechanical movement of the microscope stages are not more accurate than 1 μm, some beam-image shift is unavoidable. We have found through numerous imaging sessions that with properly aligned beam shift and beam tilt pivot points, maps with FSC_0.143_ resolution better than 3 Å can be routinely obtained.

We show here experimentally that FSC_0.143_ resolution from images taken at significant beam tilts are worse than untilted images as expected, but the effect was much smaller than predicted from the theory of a single image. Even the positions of the weaker water peaks were well preserved in this protocol. The results of Experiment 2 roughly fit (r=0.75) to a line of

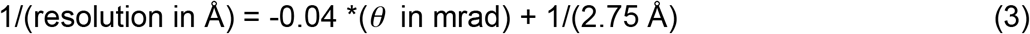

This means that a specimen with ultimate resolution of 2.75 Å can tolerate 0.76 mrad of beam tilt before its resolution is dropped below 3 Å. For a reasonably aligned microscope similar to ours, such beam tilt covers a beam-image shift of +/− 3 to 4.5 μm!

One assumption made in our explanation is that the coma-distorted particle images must align properly before averaging. The phase error is proportional to the cube of the spatial frequency. It means that distortions are very small for the low resolution component that contributes heavily to alignment. In addition, the scattering power of proteins or other macromolecules is weaker at high resolution. The square wave form represented in Figure 1A is useful for highlighting the coma effect since its structure Fourier components are equally strong across all resolution. It does not, however, represent the real world protein density that will always have some extent of information dampening at high resolution. There is further dampening at high resolution in the Fourier transform of the protein particle images that we align with SP refinement programs: counting camera MTF, and microscope optics envelope, to name a few. Taken together, the phase error coma effect should therefore not impact particle alignment very strongly. We did not observe visual differences in the quality of the 2D class averages between the low and high beam-tilt data set.

Our highest resolution reconstruction of the T20S proteasome was a 2.6 Å map taken at ~1 Å/pixel using standard processing protocols as described in the methods section but using more particles in the average, By further reducing the pixel size, we recently obtained a 2.5 Å proteasome reconstruction using the same imaging methods. This is likely because 2.5Å is at 85% of Nyquist for 1.06 Å pixel and the signal filtered through the detector quantum efficiency is too low. Our usual advice to users seeking higher resolution is to switch to smaller pixel size albeit at the cost of time.

We also would like to advise users who plan to use the beam-image shift method to first check the amount of beam tilt induced by beam-image shift on their microscope. We measured a variation from 0.17 to 0.25 mrad/μm beam-image shift across three standard Krios Titan columns. One Thermo Fisher Scientific Talos Artica column was measured at 0.47 mrad/μm (personal communication, Gabriel Lander). This value, in combination with stronger coma phase error effects at 200 kV electron voltage, made it necessary to use stage shift alone for pursuing better than 3 Å resolution SP reconstruction on the Talos Arctica (Herzik et al., 2017).

We so far have no physical explanation for the beam tilt dependency of the achieved SP resolution, which appears to roughly follow a straight line. Integration of particles of all in-plane directions suggests that the cos *Δφ* dampening of a given structure factor follows a sinc function but FSC does not directly reflect this. A full explanation will have to await more experiments or a better theory.

## Conclusions

We show that SP resolution, judged either by FSC_0.143_ or the presence of water density peaks, is not limited by the π/4 coma phase error. The averaging of large numbers of particle images allows the 3D reconstruction to reach higher resolution than theory would predict. This phenomenon makes it possible to safely use beam-image shifts as a targeting protocol when pursuing maps in the 3 Å resolution range, with a typically 2-3 times improvement in the data collection efficiency.

## Acknowledgement

Some of this work was performed at the Simons Electron Microscopy Center and National Resource for Automated Molecular Microscopy located at the New York Structural Biology Center, supported by grants from the Simons Foundation (SF349247), NyStAR, and the NIH National Institute of General Medical Sciences (Gm103310, 0D019994) with additional support from the Agouron Institute (F00316).

